## Supplementary Information for "Molecular insights into phosphoethanolamine cellulose formation and secretion"

Supplementary Figures S1-S10

Table S1

### Supplemental Figures and Captions

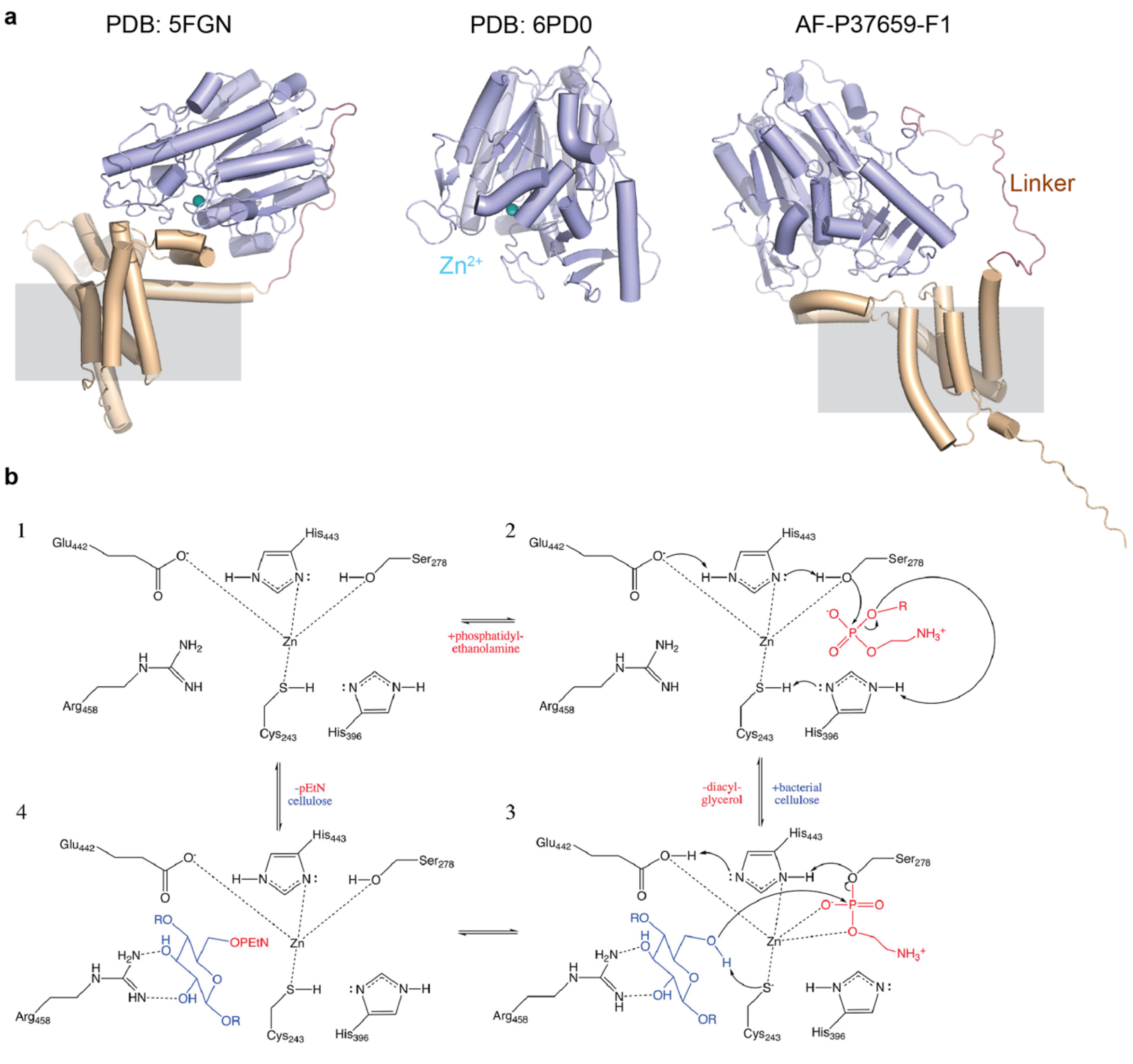


https://creativecommons.org/licenses/by-nc-nd/4.0/

**Fig. S1: BcsG is a phosphoethanolamine transferase. a**, Comparison of the crystal structures of EptA (PDB: 5FGN) and the catalytic domain of BcsG (PDB: 6PD0) with the AlphaFold-2 predicted full-length structure of BcsG. **b**, Proposed BcsG reaction mechanism as presented in reference [25].

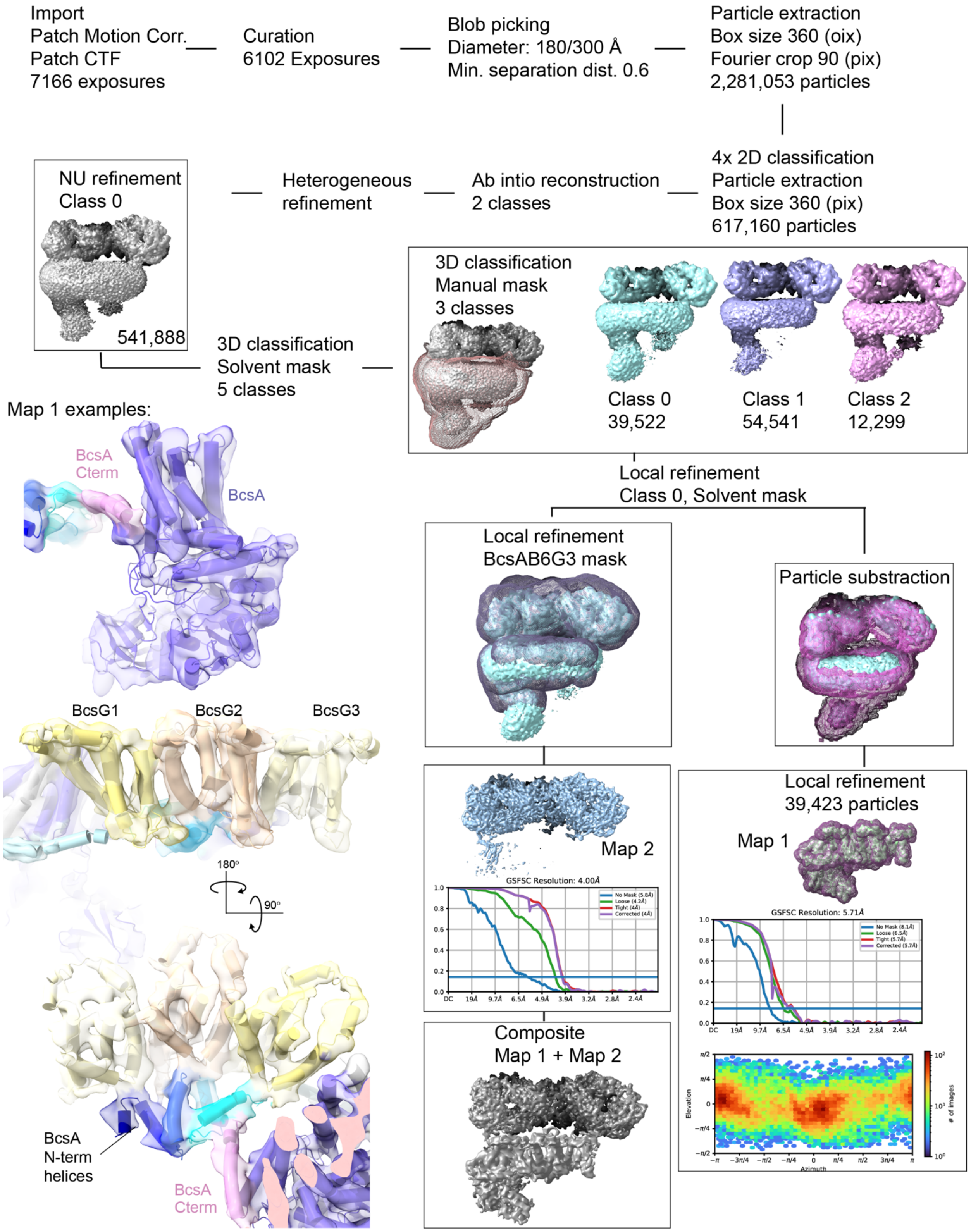


**Fig. S2: Workflow of cryo-EM data processing of the Bcs complex.** Revised processing of entry EMD-23267.

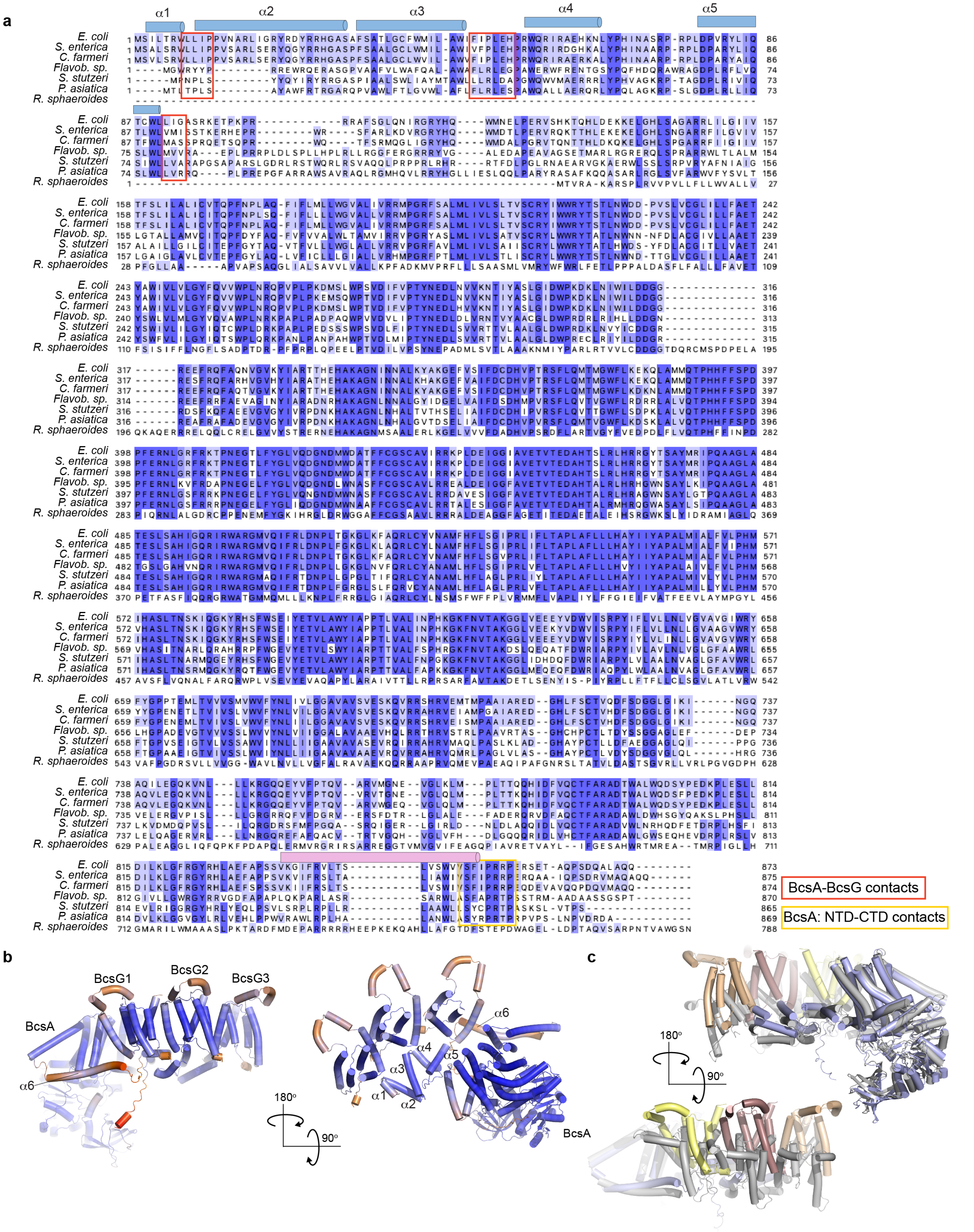


**Fig. S3: Sequence alignment of BcsA and its interaction with BcsG. a**, Sequences were aligned in MUSCLE[43] and displayed in Jalview[44]. N- and C-terminal helical elements are shown as cylinders in blue and pink, respectively. **b**, AlphaFold2-predicted complex of BcsA with three copies of

BcsG's TM segment, colored based on pLDDT score from blue to red for high to low confidence scores. **c**, Overlay of the generated AlphaFold2 predicted complex of the Ec BcsA-BcsG3 complex (colored blue for BcsA and yellow, violet and wheat for the BcsG subunits) and the model obtained after rigid body docking of the subunits into the experimental cryo EM map (colored gray).

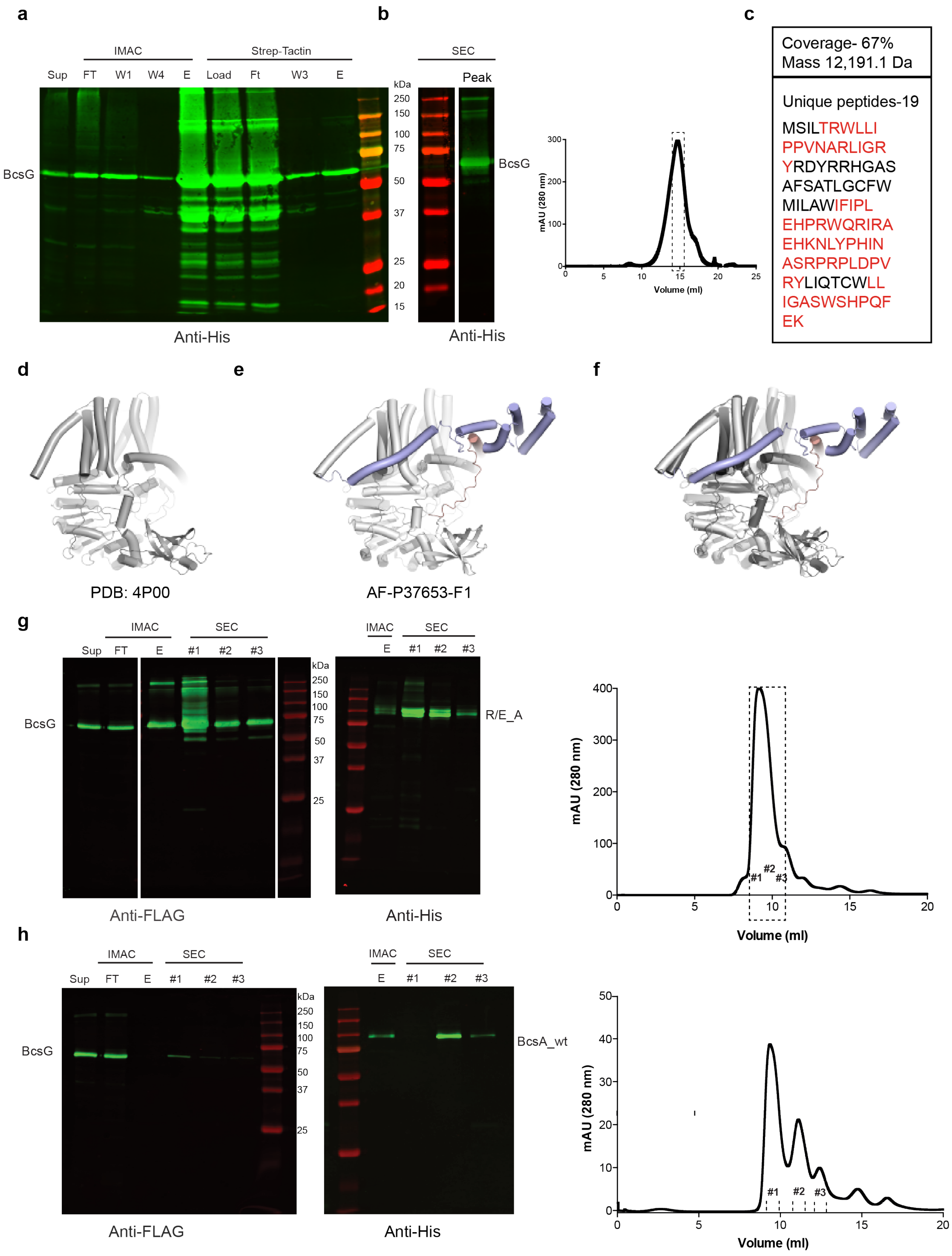


**Fig. S4: Interactions of BcsA and BcsG. a**, Anti-His Western blot of samples obtained from a tandem purification using IMAC and Strep-Tactin affinity matrices. BcsG is His-tagged and BcsA's NTD is Strep-tagged. Sup: Supernatant, FT: flow through, W: Wash step, E: Elution. **b**, Anti-His Western blot of the peak fraction after size exclusion chromatography (SEC). Right panel: Superose 6 increase SEC profile of the co-purified BcsA NTD-BcsG complex. **c**, Peptide coverage (red) of BcsA's NTD after mass spectrometry fingerprinting of a gel slice excised from an acrylamide gel at the expected molecular weight. **d** and **e**, Comparison of *Rs* (PDB: 4P00) and Ec BcsA (AF_P37653-F1). **f**, Superposition of *Rs* BcsA (PDB: 4P00) with Ec BcsA (AF_P37653-F1) by secondary structure matching. Regions absent in *Rs* BcsA are shown in blue (*Ec* BcsA NTD) and red (*Ec* BcsA C- terminus). **g**, Western blot detecting FLAG-tagged BcsG and the His-tagged chimeric BcsA after IMAC and SEC for the purified chimeric BcsAB-BcsG complex. Right panel: Supedex200 increase SEC profile**.** Sup: Supernatant, FT: flow through, E: Elution, #1-3: SEC fractions. **h**, Western blot detecting FLAG-tagged BcsG and His-tagged BcsA, respectively, of samples obtained after IMAC and SEC for the wild type BcsAB complex co-expressed with *Ec* BcsG and BcsF. Right panel: Supedex200 increase SEC profile. Solubilized membranes (Sup) have been loaded as reference for BcsG expression in the Anti-FLAG blot.

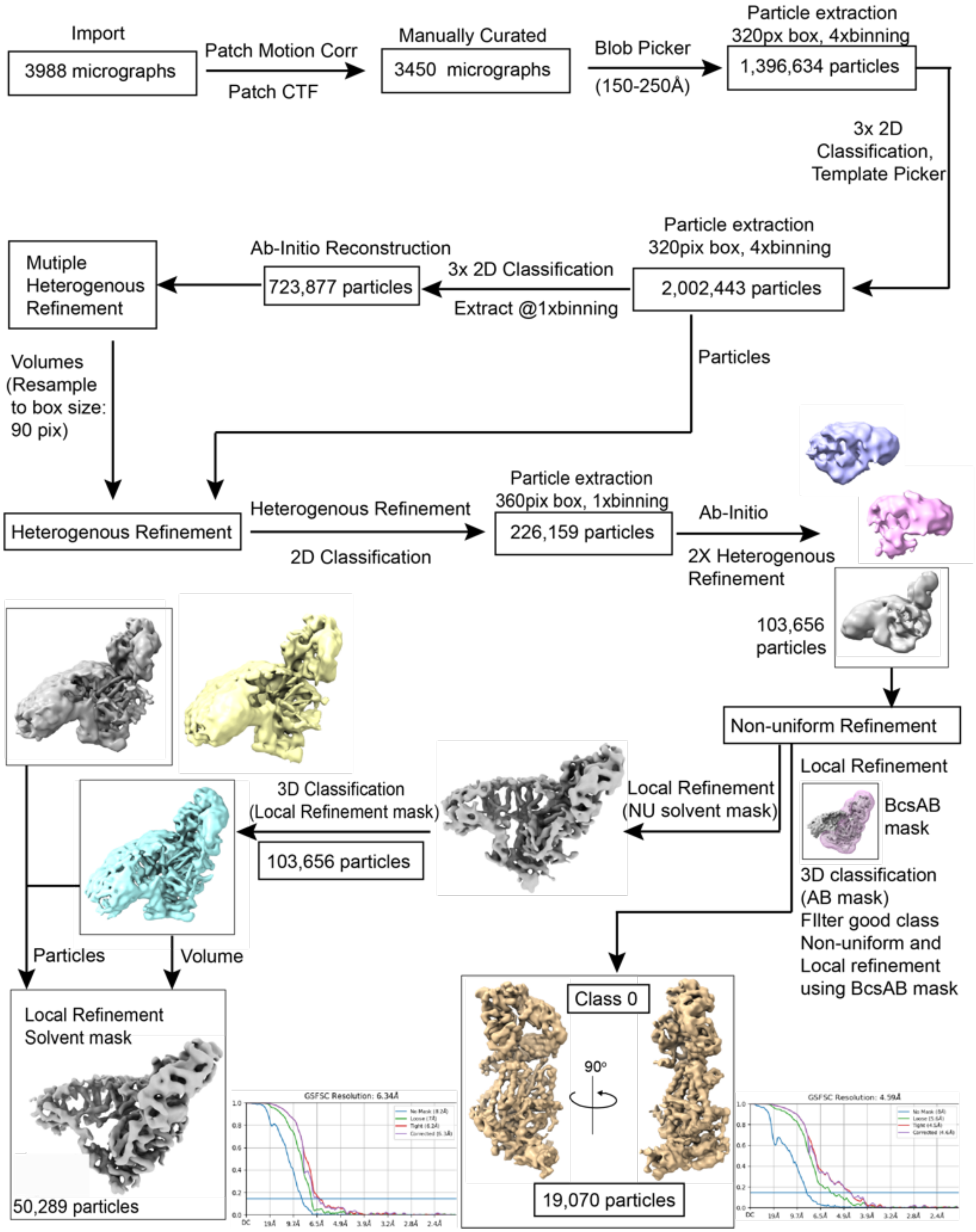


**Fig. S5: Workflow of cryo-EM data processing of the chimeric BcsAB-BcsG complex.**

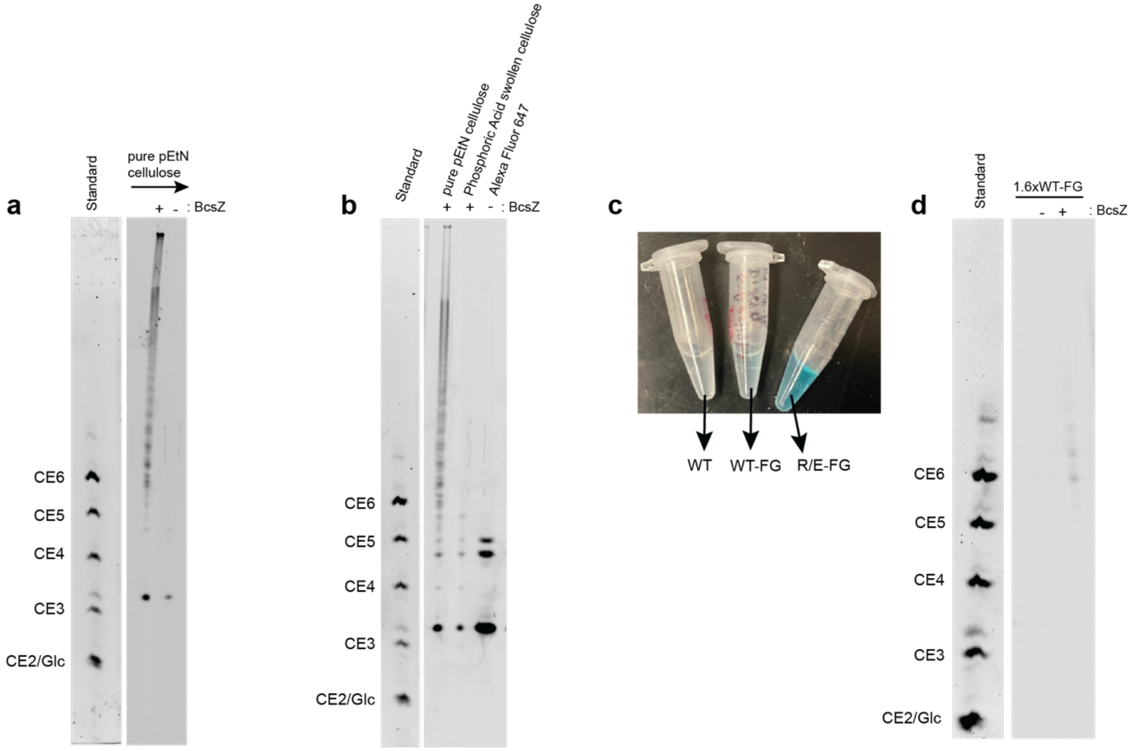


**Fig. S6: Detection of pEtN-cellulose by polysaccharide carbohydrate gel electrophoresis (PACE)**. **a**, Control labeling of purified pEtN-cellulose produced by *Ec* with Alexa Fluor 647 NHS ester followed by cellulase digestion (BcsZ). **b**, Similar as for panel (a) but including digestion of phosphoric acid swollen cellulose and an Alexa Fluor 647 NHS ester-only sample. **c**, Supernatants containing the released soluble cello-oligosaccharides after cellulase (BcsZ) digestion as described in Methods. **d**, As for panels a and b, PACE analysis of cellulose produced by IMVs containing the WT BcsAB complex co-expressed with BcsG and BcsF. 1.6-times more IMVs were used for cellulose synthesis reaction as compared to Fig. 2h. Standard: Mixture of ANTS labeled cello-oligosaccharides (DP 2 to 6) and glucose.

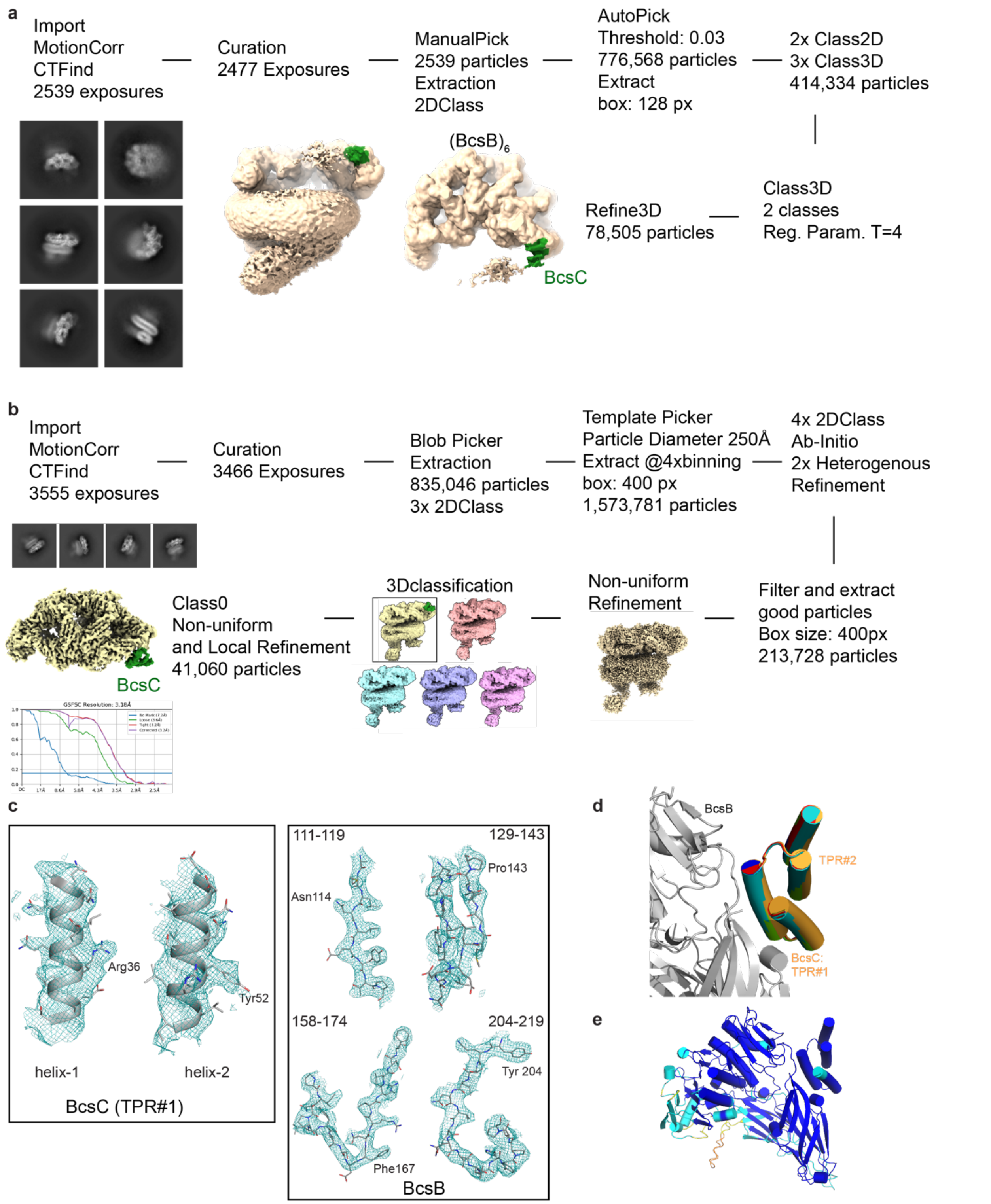


**Fig. S7: Interaction of BcsB and BcsC.** **a**, RELION Workflow of cryo-EM data processing of the Bcs complex in the presence of BcsC's periplasmic domain. **b**, CryoSPARC workflow of cryo-EM data processing of the Bcs complex containing BcsC's N-terminal two TPRs fused to BcsB. **C**, Examples of the map quality shown for TPR#1 as well region of BcsB that interacts with TPR#1, contoured at $\sigma$=5 for BcsC and $\sigma$=7 for BcsB. **d**, Five predicted AlphaFold2 models of BcsB in complex with

BcsC’s N-terminal two TPRs. BcsB for one of the models is shown as grey cartoon and BcsC TPR helices represented as cylinders are shown in different colors for models 1-5. **e**, As for panel c, but colored based on pLDDT score (blue to red for high to low confidence).

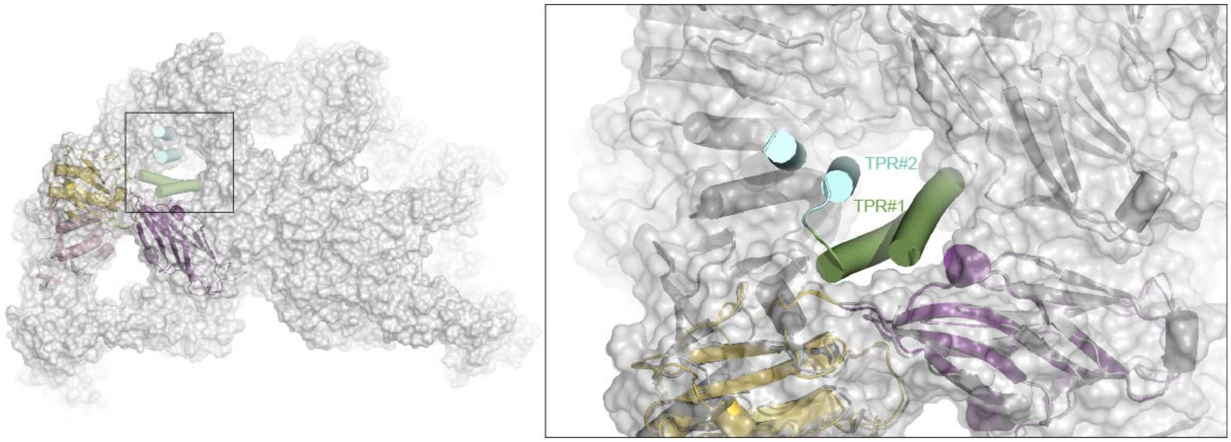


**Fig. S8 Steric constraints of BcsB and BcsC interaction.** The terminal sixth copy of BcsB in complex with BcsC was aligned with the second protomer of the BcsB semicircle to highlight steric clashes.

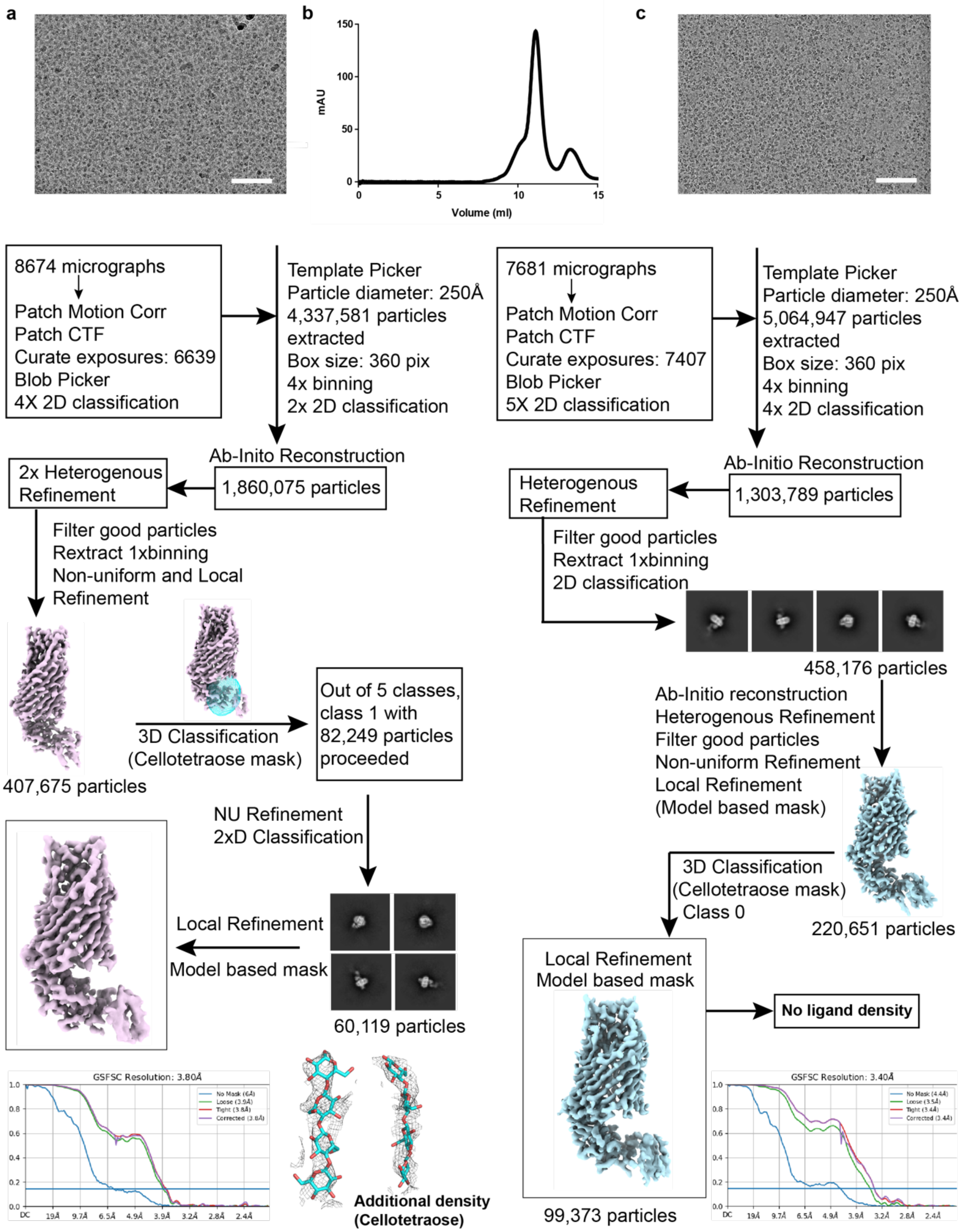


**Fig. S9 Workflow of cryo-EM data processing of BcsC with and without cellotetraose. a**, Workflow in the presence of cellotetraose with ligand density contoured at 11 σ and shown as black mesh. **b**, Gel filtration chromatogram of nanodisc-reconstituted BcsC. **c**, Workflow in the absence of cellotetraose. Scale bar on the micrograph corresponds to 100 nm.

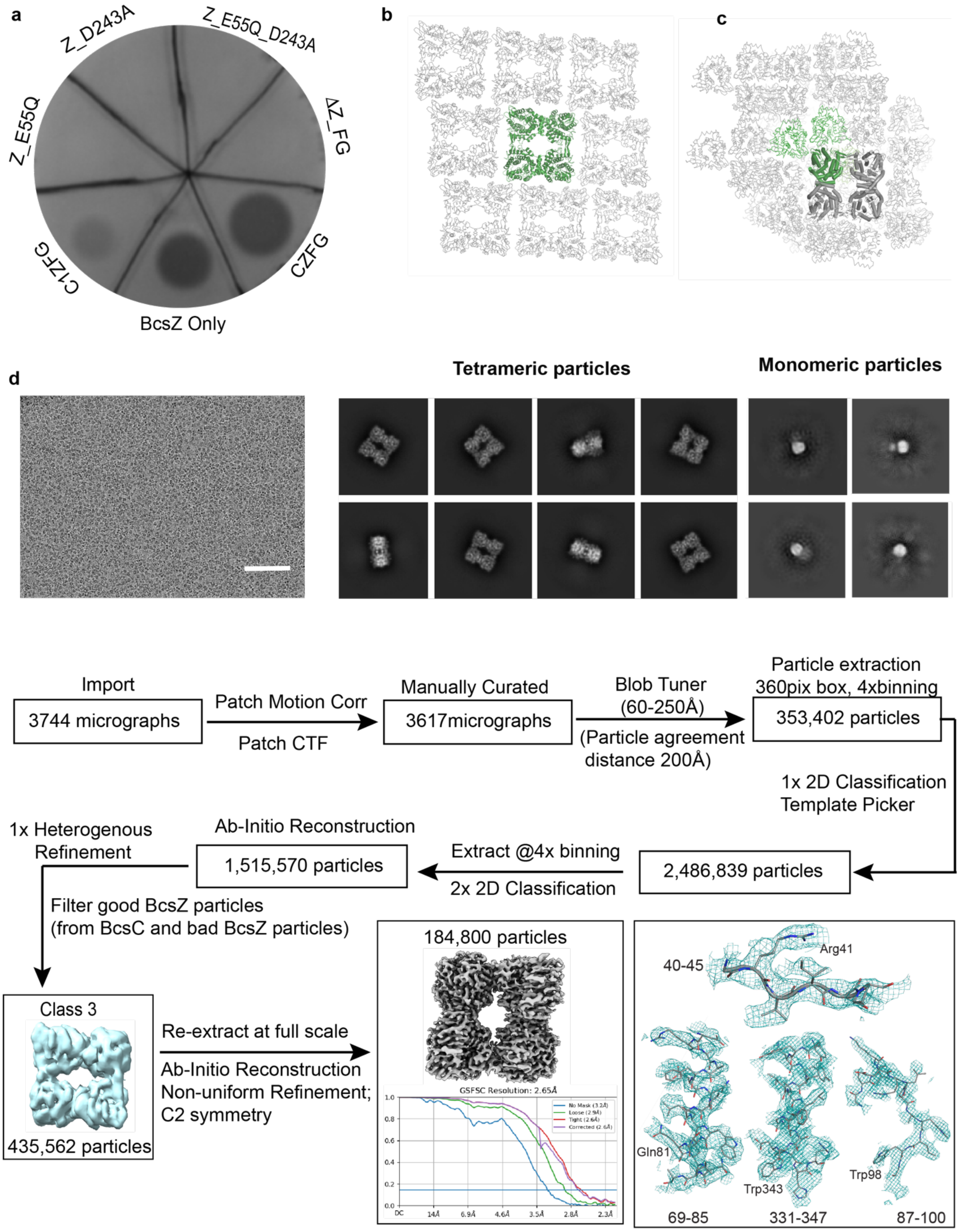
a
Z_D243A
Z_E55Q_D243A
Z_E55Q
ΔZ_FG
C1ZFG
CZFG
BcsZ Only
b
c
d
Tetrameric particles
Monomeric particles
Import
3744 micrographs
Patch Motion Corr
Patch CTF
Manually Curated
3617micrographs
Blob Tuner
(60-250Å)
(Particle agreement
distance 200Å)
Particle extraction
360pix box, 4xbinning
353,402 particles
1x 2D Classification
Template Picker
2,486,839 particles
Extract @4x binning
2x 2D Classification
Ab-Initio Reconstruction
1,515,570 particles
1x Heterogenous
Refinement
Filter good BcsZ particles
(from BcsC and bad BcsZ particles)
Class 3
435,562 particles
Re-extract at full scale
Ab-Initio Reconstruction
Non-uniform Refinement;
C2 symmetry
184,800 particles
GSFSC Resolution: 2.65Å
40-45
Arg41
Gln81
Trp343
Trp98
69-85
331-347
87-100

**Fig. S10 Function and structure of BcsZ**. **a**, Cellulase activity of overnight *Ec* cultures expressing the indicated BcsZ variants along with the full length BcsC, BcsF and BcsG subunits. ΔBcsZ: no BcsZ. BcsZ Only- indicates *Ec* cells expressing only WT BcsZ, CZFG indicates cells expressing the full-length BcsC, BcsZ, BcsF and BcsG subunits. C1ZFG: As for CZFG but expressing a BcsC construct lacking the TPRs. Cells were spotted on carboxymethylcellulose-containing agar plates, incubated at 37 °C for 48 hours and stained with CR as detailed in Methods. **b** and **c**, Crystal packing of BcsZ in space group P21 and P1, PDB entries 3QXQ and 3QXF. **d**, Workflow of cryo-EM data processing of BcsZ including a representative micrograph and 2D class averages**.** Scale bar on the micrograph: 100 nm. Examples of EM map quality, contoured at $\sigma$=7.

### Table S1: Cryo-EM data statistics and refinement.

| | Ec BcsAB6-BcsG3 (EMD-xxx) (PDB xxx) | BcsA-chimera-BcsB-BcsG complex | Ec Bcs complex with BcsC (#TPR1-18) | BcsB-BcsC fusion Ec complex (EMD-xxxx) (PDB xxxx) | BcsC Apo | BcsC Cellotetraose bound (EMD-xxxx) (PDB xxxx) | BcsZ (EMD-xxxx) (PDB xxxx) |
|---|---|---|---|---|---|---|---|
| **Data collection and processing** | | | | | | | |
| Magnification | 81,000x | 81,000x | 81,000x | 81,000x | 81,000x | 81,000x | 81,000x |
| Voltage (kV) | 300 | 300 | 300 | 300 | 300 | 300 | 300 |
| Electron exposure (e–/Å²) | 51 | 50 | 50 | 50 | 50 | 50 | 50 |
| Defocus range (μm) | -2.5 to -1 | -2 to -1 | -1.8 to -0.8 | -2 to -1 | -2.2 to -1.2 | -2.4 to -1.2 | -2.2 to -1.0 |
| Pixel size (Å) | 1.08 | 1.08 | 1.08 | 1.08 | 1.08 | 1.08 | 1.08 |
| Symmetry imposed | C1 | C1 | C1 | C1 | C1 | C1 | C2 |
| Initial particle images (no.) | 2,281,053 | 2,002,443 | 776,568 | 1,573,781 | 5,064,947 | 4,337,581 | 2,111,486 |
| Final particle images (no.) | 39,522 | 50,289 | 78,505 | 41,060 | 99,373 | 60,119 | 184,800 |
| Map resolution (Å) | 5.3 | 6.3 | 12 | 3.2 | 3.4 | 3.8 | 2.7 |
| FSC threshold | 0.143 | 0.143 | 0.143 | 0.143 | 0.143 | 0.143 | 0.143 |
| Map resolution range (Å) | 5.3-100 | 2.2-98.8 | | 2.35-51.6 | | 3.1-56.3 | 2.4-39 |
| **Refinement** | | | | | | | |
| Initial model used (PDB code) | AlphaFold2 and 7L2Z | AlphaFold2 | AlphaFold2 | AlphaFold2 | AlphaFold2 | AlphaFold2 | 3QXQ |
| Model resolution (Å) | 5.7 | | | 3.2 | 3.4 | 3.8 | 2.60 |
| FSC threshold=0.143 | | | | | | | |
| Map sharpening *B* factor (Å²) | 347 | | | -58.7 | -65.9 | -78.3 | -80.1 |
| Model composition | | | | | | | |
| Non-hydrogen atoms | 70365 | | | 5610 | 4320 | 4641 | 10920 |
| Protein residues | 5179 | | | 719 | 554 | 587 | 1352 |
| Ligands | 0 | | | 0 | 0 | CE5: 1 | 0 |
| *B* factors (Å²) | Rigid body | | | | | | |
| Protein | docked | | | 119.68 | 69.24 | 142.74 | 13.40 |
| Ligand | only | | | - | | 184.95 | - |
| R.m.s. deviations | | | | | | | |
| Bond lengths (Å) | | | | 0.002 | 0.002 | 0.003 | 0.003 |
| Bond angles (°) | | | | 0.512 | 0.433 | 0.664 | 0.677 |
| Validation | | | | | | | |
| MolProbity score | | | | 1.49 | 1.39 | 1.79 | 1.46 |
| Clashscore | | | | 4.74 | 5.72 | 8.08 | 8.42 |
| Poor rotamers (%) | | | | 1.28 | 0.89 | 0 | 0.71 |
| Ramachandran plot | | | | | | | |
| Favored (%) | | | | 97.05 | 97.64 | 94.87 | 99.429 |
| Allowed (%) | | | | 2.95 | 2.36 | 5.13 | 0.6 |
| Disallowed (%) | | | | 0 | 0 | 0 | 0 |
